# Climate explains recent population divergence, introgression and persistence in tropical mountains: phylogenomic evidence from Atlantic Forest warbling finches

**DOI:** 10.1101/439265

**Authors:** Fábio Raposo do Amaral, Diego F. Alvarado-Serrano, Marcos Maldonado-Coelho, Katia C. M. Pellegrino, Cristina Y. Miyaki, Julia A. C. Montesanti, Matheus S. Lima-Ribeiro, Michael J. Hickerson, Gregory Thom

## Abstract

Taxa with disjunct distributions are common in montane biotas and offer excellent opportunities to investigate historical processes underlying genetic and phenotypic divergence. In this context, subgenomic datasets offer novel opportunities to explore historical demography in detail, which is key to better understand the origins and maintenance of diversity in montane regions. Here we used a large ultraconserved elements dataset to get insights into the main biogeographic processes driving the evolution of the Montane Atlantic Forest biota. Specifically, we studied two species of warbling finches disjunctly distributed across a region of complex geological and environmental history. We found that a scenario of three genetically differentiated populations is best supported by genomic clustering methods. Also, demographic simulations support simultaneous isolation of these populations at ~10 kya, relatively stable population sizes over recent time, and recent gene flow. Our results suggest a dual role of climate: population divergence, mediated by isolation in mountain tops during warm periods, as well as population maintenance - allowing persistence mediated by shifts in elevation distribution during periods of climate change, with episodic bouts contact and gene flow. Additional support for the role of climate comes from evidence of their contact in a recent past. We propose that two major gaps, which we call São Paulo and Caparaó subtropical gaps, have been historically important in the divergence of cold adapted organisms in the Atlantic Forest, and could be associated to cryptic diversity. Finally, our results suggest that shallow divergence and past gene flow may be common in montane organisms, but complex demographic histories may be detectable only when using subgenomic or genomic datasets.

## Introduction

Montane organisms often exhibit intriguing patterns of disjunct distributions, which are frequently associated to phenotypic divergence. Among the many mechanisms that have been proposed to explain such patterns, the ones with the strongest support are the historical fragmentation of formerly continuous habitats and dispersal (Mayr & Diamond 1976; Patton & Smith 1992; Knowles & Massatti 2017) asssociated with rugged topography and climate fluctuations. That is, changes in climate over time, which are well known to promote shifts in the elevational distributions of suitable habitats and associated organisms (Hooghiemstra & Van der Hammen 2004, Moritz et al. 2008, Chen et al. 2009), could lead to cyclical isolation and connection of population through temporal bridges (Brown 1971, McCormack et al. 2009). During the late Pleistocene glacial periods, warmer periods often lead to isolation in high-elevation habitat pockets, whereas colder periods often lead to range expansion into lower elevations and previously unsuitable regions. Under this scenario, warmer periods (such as the current interglacial) would create opportunities for population divergence by genetic drift and natural selection, while colder periods might promote secondary contact and genetic homogenization of previously isolated populations. This interplay between selection and drift with topography and a dynamic climate could lead not only to genetic divergence, but to the evolution of reproductive isolation, thus rendering mountains as potential hotspots of population divergence and speciation (Fjeldsa 1994, McCormack et al. 2009). Alternatively, isolation could be initially mediated by topographic changes. For example, the splitting of formerly continuous highlands into multiple isolated mountain ranges could be the result of geological changes, and organisms adapted to higher habitats would then become separated by ecologically unsuitable valleys. Under such scenario, population divergence should be temporally associated to geotectonic changes (Badgley 2010).

These biogeographic processes likely have played an important role in the building up of the enormous diversity found in mountain regions (Graham et al. 2014, Antonelli 2015; Knowles & Massatti 2016). However, their exact contribution is not well understood. In particular, the evolutionary processes underlying the high levels of biological diversity in the South American Montane Atlantic Forest are yet to be elucidated. The Montane Atlantic Forest (hereafter MAF) is a cradle of biodiversity (Stotz et al. 1996) and has great potential to provide a wealthy natural laboratory for evolutionary research since its high levels of endemism likely encompass a plethora of diversification mechanisms. In particular, MAF organisms showing phenotypic breaks that coincide with major highlands are especially interesting models for montane phylogeography, as such pattern presumably reflects genome-wide divergence resulting from an interplay between climate, topography and the evolutionary outcomes of drift, selection and gene flow. Of these, vagile organisms such as birds may be particularly useful models to understand large-scale historical events, as their fidelity to specific habitats and high dispersal ability may promote genetic differentiation and homogenization, except in face of strong events of habitat isolation.

Much of what is known about the Atlantic Forest comes from studies of either lowland species or species with broad elevational distributions (e.g. Cabanne et al. 2007, Carnaval et al. 2009. Maldonado-Coelho 2012, Amaral et al. 2013, Carnaval et al. 2014), and less than a handful of phylogeographic studies have investigated cold-adapted species whose distributions include mostly montane habitats (e.g. Amaro et al. 2013, Batalha-Filho et al. 2012, Peres et al. 2015, Frikowski et al. 2016, Françoso et al. 2016, Pie et al. 2018). Conclusions from the few studies on MAF-inhabiting species are often discordant; consequently, there is still little consensus about what are the major drivers of diversification in the MAF. A common finding by many of them is the lack of both population structure and strong demographic fluctuations, which could speak against Pleistocene climatic fluctuations as major drivers of population divergence and diversification in MAF organisms. Importantly, all but one (Pie et al. 2018) of the studies performed so far were based on five or less loci, raising the possibility that shallow population structure—which is expected under recent climate fluctuations—may have been overlooked (Amaral et al. 2018). Furthermore, none of these studies have performed demographic analyses explicitly testing for historical gene flow among currently isolated and genetically structured populations—a prediction of montane climate-driven dynamics.

Here we use the largest population-level sub-genomic dataset of any MAF organism and coalescent-based methods to ask (1) how does genetic variation relates to distribution breaks in the MAF, and (2) what are the influences of climate and topographic evolution (i.e. mountain building) on the current patterns of genetic variation in endemic Atlantic Forest montane birds. To address these questions we use two common cold-adapted species of Atlantic Forest warbling finches as models: the Gray-Throated Warbling Finch (*Microspingus cabanisi*) and the Buff-Throated Warbling Finch (*Microspingus lateralis*). Their distributions and phenotypic breaks are congruent with major isolated highlands in S and SE Brazil (Assis et al. 2007), and a detailed phylogeographic analysis of these species represents a fundamental step towards a better understanding of population history in MAF and biotic diversification in tropical montane systems as a whole.

## Methods

### Study system: biological models and geological and environmental history of the MAF

The two warbling finches studied here are sister-species (Amaral *et al.* 2015), once considered conspecific (Assis et al. 2007), and whose combined distributions cover most of the MAF. The northern species, *M. lateralis* inhabits highland areas above 900 m of two major isolated mountain ranges in SE Brazil, the Mantiqueira and the northern portion of the Serra do Mar. This species also occurs in the Caparaó highlands, an isolated northernmost massif of the Mantiqueira mountain range (Fig.1). The southern species, *M. cabanisi*, occurs in the southern portion of Serra do Mar and also in the Serra Geral mountain range mostly above 900 m, but also in lower elevation areas towards its southern limit, which includes areas in Argentina and Paraguay (Assis et al. 2007). These two species are currently separated by more than 100 km of lower elevation forest habitats (Fig. 1). They differ phenotypically in both plumage and vocalizations. Plumage differences are mostly in ventral color (yellow in *M. lateralis* and gray in *M. cabanisi*) and amount of white in tail feathers (more extensive in *M. cabanisi*), whereas the songs and the calls of these species differ in syntax and harmonic structure, respectively (Assis et al 2007).

**Figure 1.**
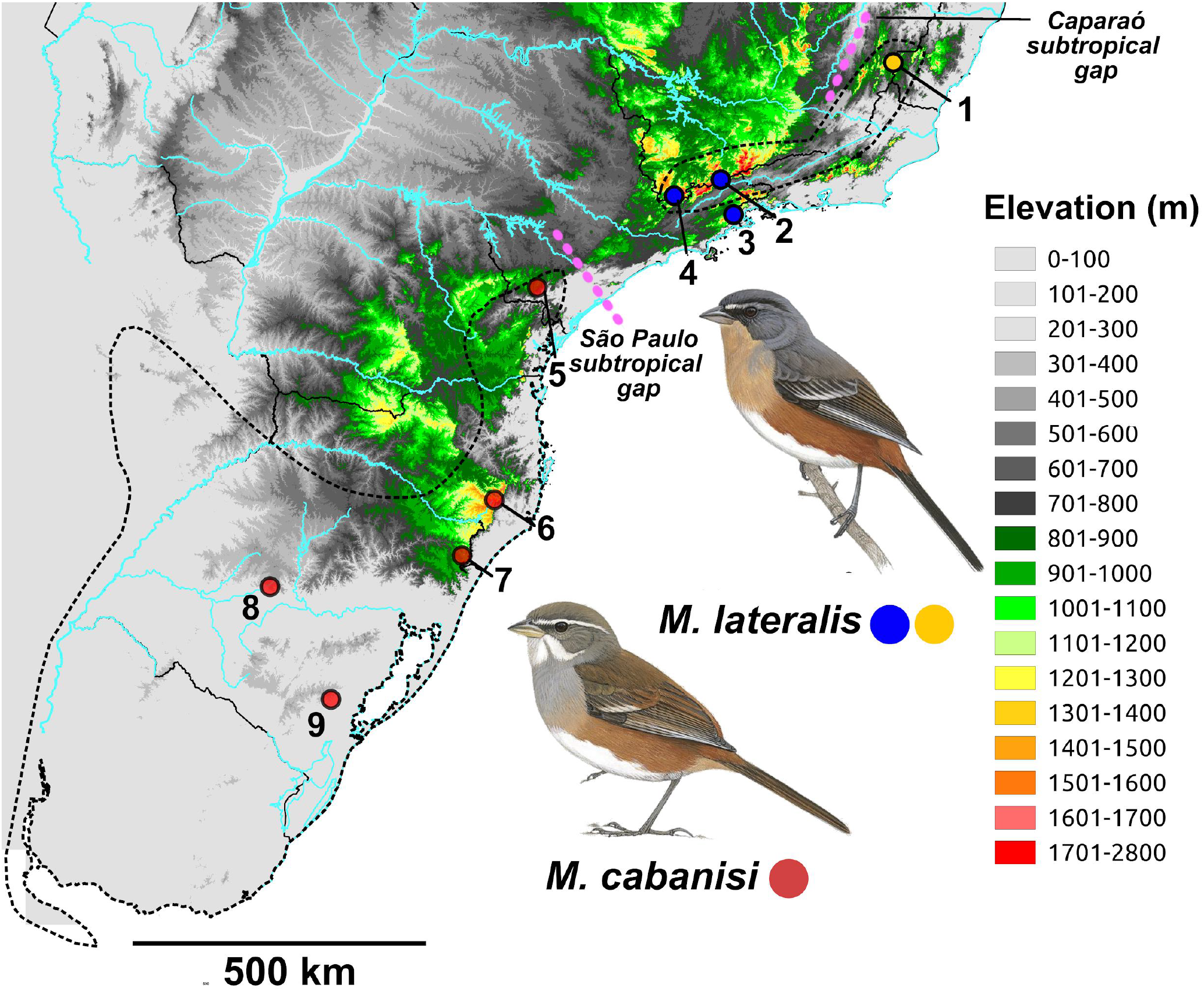
Localities sampled. Blue and yellow circles represent *Microspingus lateralis* while red circles represent *Microspingus cabanisi*. Major highlands include Caparaó (1), Northern Serra do Mar (2), Mantiqueira (3 and 4), Southern Serra do Mar (5) and Serra Geral (6 and 7). Two lower elevation regions that mark phenotypic and genotypic shifts, named here Caparaó and São Paulo subtropical gaps, are indicated by pink dashed lines. See Supplementary Information Table S1 for details on samples and coordinates.

The tectonic processes that originated the eastern Brazilian highlands as well as their ages are still a matter of debate. In one hand, the major uplift events were suggested to have started during the lower Cretaceous (ca. 120 mya) as a consequence of the Gondwanaland breakup (Melo et al. 1985, Franco-Magalhães et al. 2014). On the other hand, the uplift is proposed to have resulted from the three main pulses of Andean orogeny, with the early ages of the mountain ranges (i.e. Serra do Mar) estimated to be ca. 90 mya (Meisling et al. 2001, Karl et al. 2013). Regardless, a range of evidence indicates subsequent multi-episodic uplift events during the upper Cretaceous, the Tertiary and the Quaternary (Freitas 1951, Melo et al. 1985, Almeida & Carneiro 1998, Modenesi-Gauttieri et al. 2002, Tello Saens et al. 2003, Hackspacher et al. 2004, Franco-Magalhães et al. 2014), with the uplift of the Serra do Mar mountain range thought to have extended until recent times (i.e. Holocene, Cogné et al. 2012).

Available evidence from fossil pollen records in southern and southeastern Brazil highland sampling sites (> 900 m a.s.l.) at Mantiqueira, Serra do Mar and Serra Geral mountain ranges suggest broadly concordant cold and dry conditions during the last glacial maximum, with forests completely absent or likely present only along ravines and/or at lower altitudes (Behling 1997, 2007, Behling & Pillar 2007, Behling et al. 2007, Oliveira et al. 2008, Ledru et al. 2009, Behling & Safford 2010, Oliveira et al. 2012). The onset of a moister climate after this period favored the return of montane forest species (Behling 1997, Behling & Pillar 2007, Behling et al. 2007, Ledru et al. 2009, Behling and Safford 2010, Oliveira et al. 2012). Importantly, the highland vegetation seems to have had a complex dynamics during the last 130,000 yr BP (i.e. late Pleistocene and Holocene), with multiple shifts in community composition and elevational distribution of montane forest plant species (Ledru et al. 2009, Behling & Safford 2010, Oliveira et al. 2012).

### Sampling and laboratory methods

We sampled 90 individuals (50 of *M. cabanisi* and 40 of *M. lateralis*) in nine localities—10 per locality, spread throughout the species’ known distributions (Fig. 1, Supplementary Information Table S1). Specimens were attracted by playback and collected as described in Amaral *et al.* 2012 (see Acknowledgments for permit number and Table S1 for specimen and tissue collection holdings).

We extracted total DNA from pectoral muscle using the Qiagen DNeasy kit (Valencia, CA) according to the manufacturer’s protocol, including a RNAse treatment. We obtained genomic data using sequence capture and Illumina sequencing of ultraconserved elements (UCEs) using standard protocols (Fairlocth *et al.* 2012) with a few modifications: enrichment was done using 650 probes, targeting 634 loci covering all *Gallus gallus* chromosomes; use of 100 bp paired-end Illumina Hiseq 2000 sequencing run, and use of 16 cycles in both pre- and post-capture PCR reactions. Sequencing was performed in two lanes. Library preparation was performed by RAPiD Genomics (Gainesville, FL, USA).

### Sequence quality control, mtDNA, UCE assembly and SNP calling

We sorted raw sequences by individual tags using Illumina’s Casava software. Initial quality control was performed using FastQC 0.10.1 (Andrews 2014). Adapters, barcodes and low quality regions were trimmed using Illumiprocessor 2.0.7 (Faircloth 2014), which processes Illumina sequences using Trimmomatic 0.32.1 (Bolger *et al.* 2014). Assembly, removal of non-UCE loci and final loci aligment was performed with Phyluce 1.4 (Faircloth 2014). The largest contig of each locus was used as a reference for mapping individual reads using BWA-mem (Li 2013) and SNP calling, which was performed using GATK (McKenna *et al.* 2010). We kept only >Q30 SNPs for downstream analyses. We sampled one random SNP per locus from the collection of all zero-missing biallelic SNPs recovered for the locus. We removed Z-linked SNPs using a local BLAST search based on the zebra-finch Z chromossome (Emsembl taeGut3.2.4) to avoid any bias due to the idiosincratic evolution of sex chromossomes and different ploidy (unfortunately the limited number of Z-linked SNPs recovered precluded us from running independent analyses on this dataset). Since mtDNA is a common subproduct of sequence capture experiments (see Amaral *et al.* 2015), we also isolated complete cythochrome b sequences from assemblies that were generated by Phyluce using a local BLAST search based on the complete mitogenome of *Microspingus lateralis* (Genbank NC_028039.1) to obtain a Median Joining Network using POPART v 1.7 (Leigh & Bryant 2015).

### Population structure

We explored population structure first with sparse non-negative matrix factorization (sNMF, Frichot et al. 2014, implemented in the R package LEA, Frichot & Olivier 2015) as well as with the multivariate Discriminant Analysis of Principal Components (Jombart *et al.* 2010). The sNMF runs were performed for a range of K values from 1 to 9 (following the number of localities), with 500 runs per K. We used the minimum cross-entropy method to identify the best fitting K. Runs with the lowest values of minimum entropy were selected. Since sNMF’s regularization parameter alpha may affect the inferences (Frichot et al. 2014), we evaluated the results under different values of alpha (1, 10, 100 and 1,000). To further assess population structure we ran a DAPC analysis using a k-clustering algorithm to determine the *a priori* grouping of individuals based on the Bayesian Information Criterion (BIC). The first set of principal components accounting for 80% of the variance was included in the analyses.

### Inference of historical processes

Based on the results from the previous analyses (Fig. 2), we tested 12 alternative models to assess the most likely diversification scenario for *Microspingus*. The models tested explored the relationship of the three genetic clusters inferred by sNMF, the presence of gene flow and instantaneous population size change after divergence (Fig. 3). To compare simulated data under each specific demographic model with the empirical data we implemented a coalescent model-based approach using Fastsimcoal2 (FSC; Excoffier et al. 2013). Since FSC summarizes the complexity of the data by using the site frequency spectrum (SFS) as summary statistics, we first estimated the empirical multi-SFS (single SFS with all populations included) in ∂a∂i 1.7 (Gutenkunst et al. 2009). For each demographic model we ran 50 independent replicates, retaining the parameters that maximized the composite likelihood across all iterations. Parameter optimization was performed through 50 cycles of the Brent algorithm and the composite likelihood calculated using 100,000 simulations per replicate. The runs with the highest likelihood of each model were used in model selection using Akaike Information Criterion (AIC; Akaike 1973), AIC=2k-2ln(L), where k is the number of parameters estimated in the model and L the composite likelihood value. The model with best fit had its confidence intervals estimated with 50 parametric bootstrap runs, by simulating multi-SFSs under the maximum likelihood estimates and re-estimating parameters for each of these simulated data sets. For all simulations, we used a rate of 2.5 × 10^-9 substitution per site per generation (average of all avian species analysed in Nadachowska-Brzyska et al. 2015) and assumed a generation time of 2.33 years (Maldonado-Coelho 2012).

**Figure 2.**
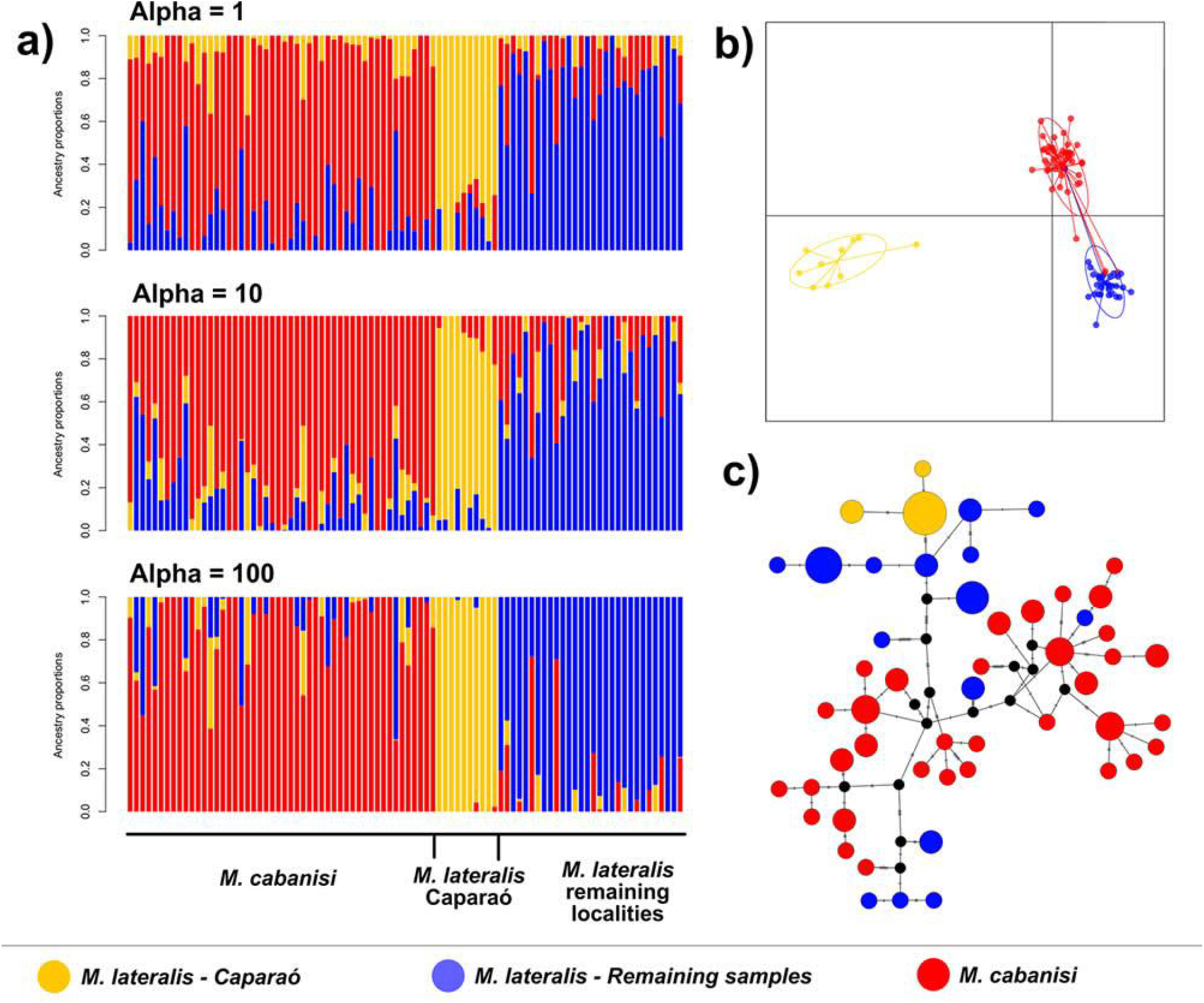
Plots depicting population structure according to (a) the best fitting sNMF model K=3 based on different values of the regularization parameter alpha (1, 10, 100) and (b) the three clusters inferred with DAPC and (c) cytochrome-b median-joining network. Black dots in the haplotype network indicate inferred haplotypes.

**Figure 3.**
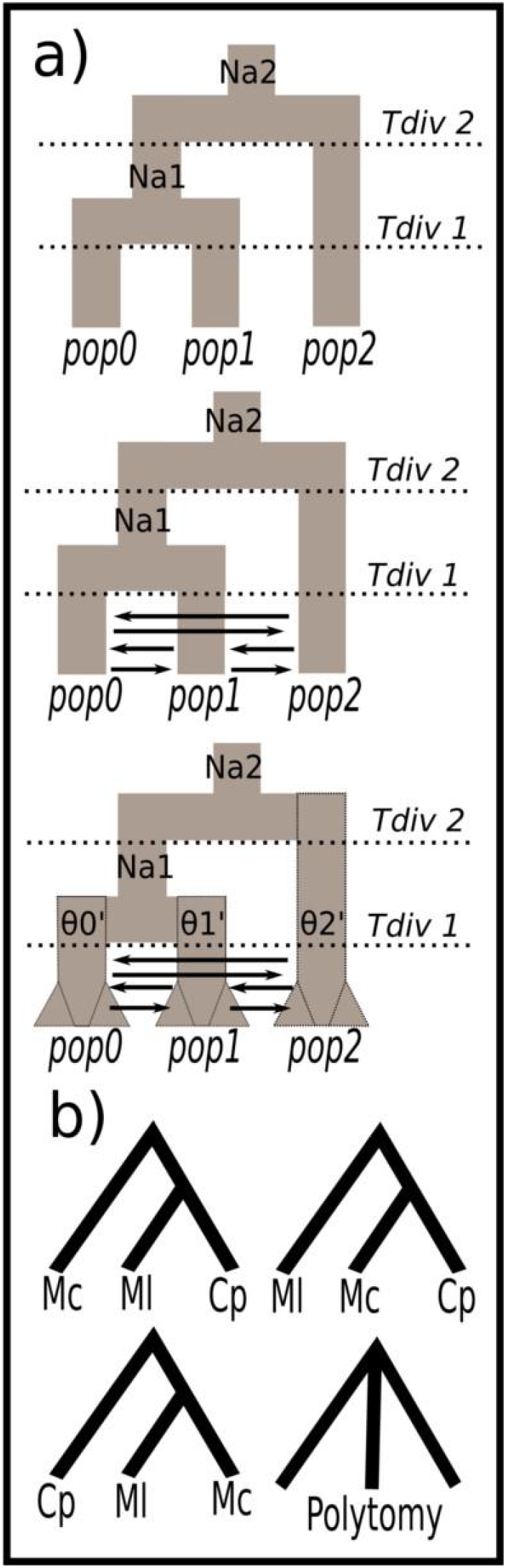
Demographic models simulated in Fastsimcoal2 for (a) three populations and (b) alternative topologies tested under these demographic scenarios, in a total of 12 simulated models. Na – ancestral effective population size; θ’ – effective population size before population size change; Tdiv – divergence time; Mc – *Microspingus cabanisi*; Ml – *Microspingus lateralis*; Cp - Caparaó population of *Microspingus lateralis*

## Results

### Sequencing results

We obtained 634 UCE loci with median length of 633 bp (range: 216-811 bp) and 8,465 SNPs passed our quality filtering. Each UCE locus had a median of 13 SNPs (range: 1-35). The final dataset containing 1 randomly sampled SNP per variable locus had 588 SNPs.

### Population structure

sNMF runs indicated three groups based on the best cross-entropy values across alpha values, with the only exception of alpha of 1,000 showing a smaller cross-entropy value for a single group (Supporting Information Figure S1). Plots based on alpha of 1, 10 and 100 for K=3 support the Caparaó population of *M. lateralis*, the remaining *M. lateralis* populations, and *M. cabanisi* as three distinct clusters with somewhat varying levels of ancestry coefficients (Fig. 2). DAPC based on k-clustering algoritm suggests two as the best number of groups, altough BIC values are very similar among K=1, K=2 and K=3 (246.7343, 246.5362 and 247.0419, respectively, Supporting Information Figure S2). These two clusters correspond to: 1) the Caparaó population of *M. lateralis* + some *M. cabanisi* individuals and 2) remainig *M. lateralis* individuals + remaining *M. cabanisi* individuals (Supporting Information Figure S2). If the three sNMF populations are considered in the DAPC analysis, the groupings are similar: one of them included only *M. lateralis* Caparaó individuals and the other two mostly matched *M. cabanisi* and *M. lateralis* excluding Caparaó (Fig. 2 and Supporting Information Fig. S1). Accordingly, the mtDNA haplotype network showed no haplotypes shared among populations, although some haplotypes of *M. lateralis* were more closely related to those of *M. cabanisi* than to conspecific haplotypes, while Caparaó haplotypes formed a cluster (Fig. 2).

### Demographic relationships and population history

The best demographic model according to AIC assumed a polytomy of the three populations, presence of assymetric gene flow and stable population sizes after the divergence (Fig. 3; Tables 1 and 2). The second and third best-fit models included alternative branching (instead of simultaneous divergence) with gene flow, but neither supported population size changes (Table 1). Parameter estimations based on the best-fit model supported current effective population sizes (Ne) that were positively related with the modern range size of the three lineages. The geographically most restricted population (i.e. Caparaó) had the smallest Ne, with ~4,000 individuals, followed by the populations with intermediate (rest of *M. lateralis*) and largest (*M. cabanisi*) geographic ranges, with respectively ~11,000 and ~19,500 individuals (Table 2). The divergence time estimate supports a very recent (Late Pleistocene) event for the simultaneous separation of the three populations at approximately 12,400 years ago (5,325 generations, Table 2).

**Table 1.**
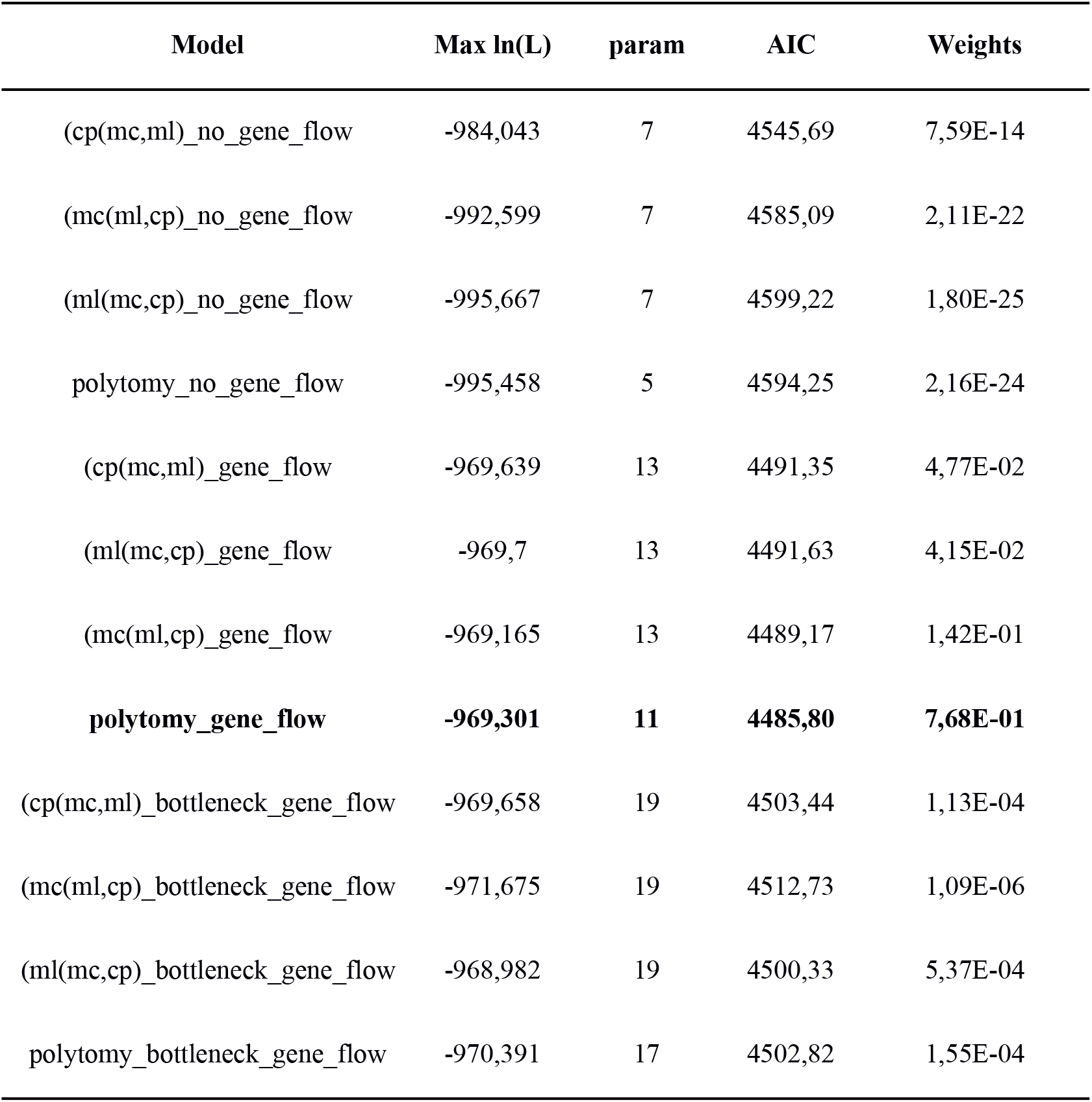
Composite likelihood (Max ln(L)), number of parameter (param), Akaike information criterion (AIC) and relative contribution (Weights) for each of the demographic models tested. The name of the models represent the topology, in newick format and the demographic syndromes tested (see Figure 3). mc - *Microspingus cabanisi*; ml - *Microspingus lateralis*; cp - Caparaó population of *Microspingus lateralis.* In bold: best model.

| Model | Max ln(L) | param | AIC | Weights |
| --- | --- | --- | --- | --- |
| (cp(mc,ml)_no_gene_flow | -984,043 | 7 | 4545,69 | 7,59E-14 |
| (mc(ml,cp)_no_gene_flow | -992,599 | 7 | 4585,09 | 2,11E-22 |
| (ml(mc,cp)_no_gene_flow | -995,667 | 7 | 4599,22 | 1,80E-25 |
| polytomy_no_gene_flow | -995,458 | 5 | 4594,25 | 2,16E-24 |
| (cp(mc,ml)_gene_flow | -969,639 | 13 | 4491,35 | 4,77E-02 |
| (ml(mc,cp)_gene_flow | -969,7 | 13 | 4491,63 | 4,15E-02 |
| (mc(ml,cp)_gene_flow | -969,165 | 13 | 4489,17 | 1,42E-01 |
| **polytomy_gene_flow** | **-969,301** | **11** | **4485,80** | **7,68E-01** |
| (cp(mc,ml)_bottleneck_gene_flow | -969,658 | 19 | 4503,44 | 1,13E-04 |
| (mc(ml,cp)_bottleneck_gene_flow | -971,675 | 19 | 4512,73 | 1,09E-06 |
| (ml(mc,cp)_bottleneck_gene_flow | -968,982 | 19 | 4500,33 | 5,37E-04 |
| polytomy_bottleneck_gene_flow | -970,391 | 17 | 4502,82 | 1,55E-04 |

**Table 2.**
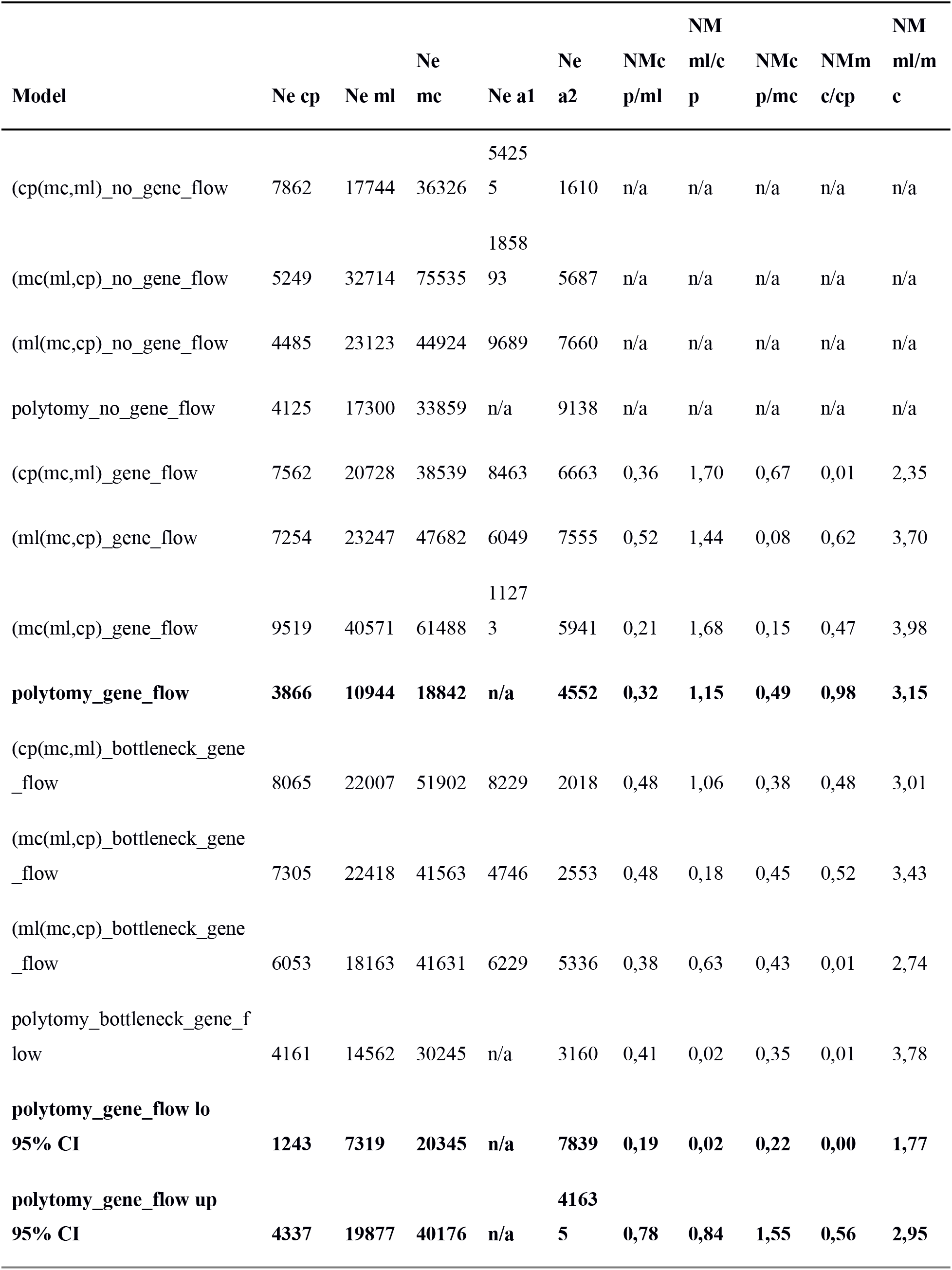
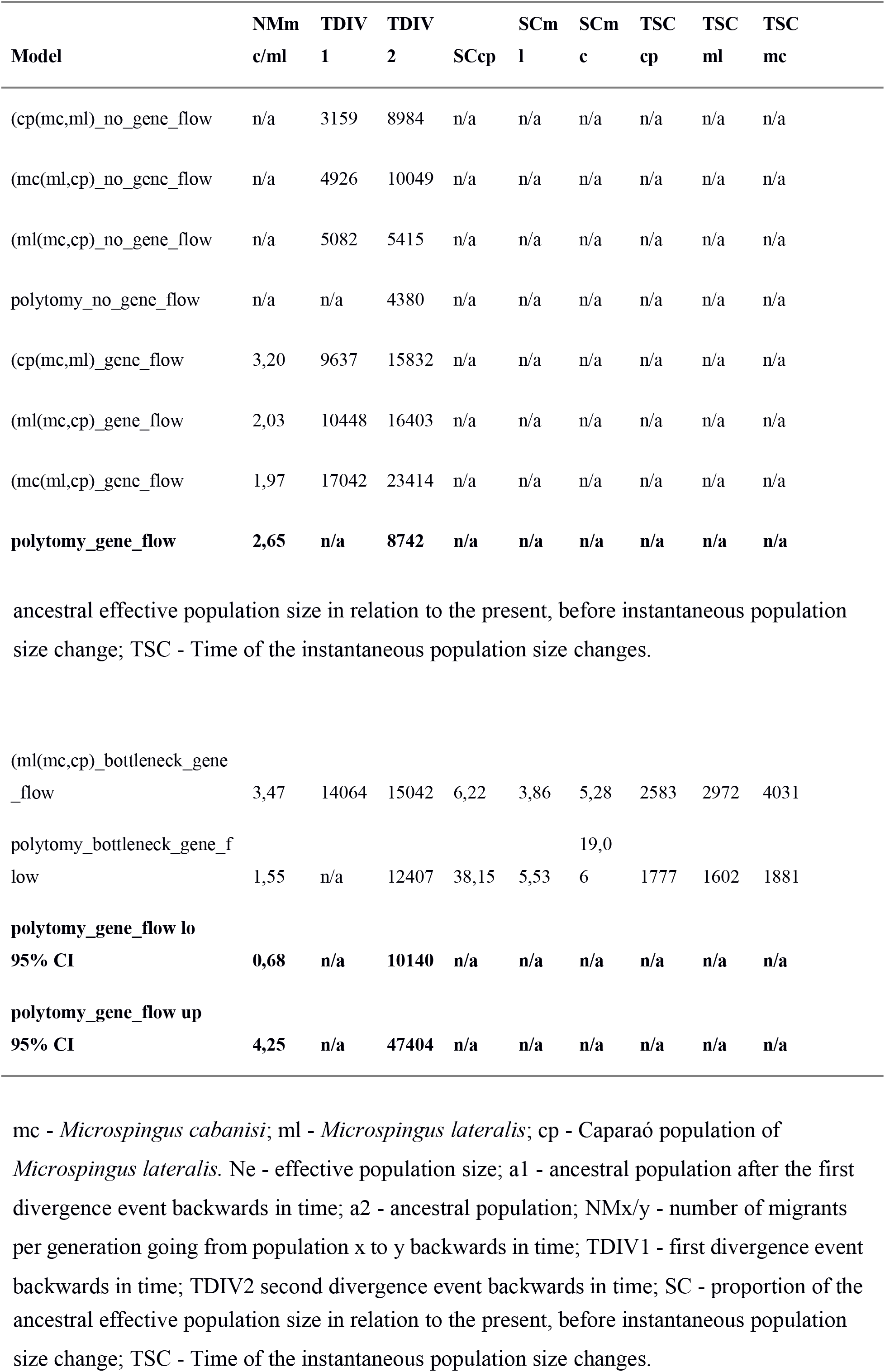
Parameter values of simulated models with the best likelihood in Fastsimcoal2. The name of the models represent the topology, in newick format and the demographic syndromes tested (see Figure 3). In bold: best model.

The estimated gene flow between *M. cabanisi* and *M. lateralis* (excluding Caparaó) was greater (3.15 and 2.65 individuals per generation from *M. lateralis* to *M. cabanisi* and from *M. cabanisi* to *M. lateralis*, respectively) than between *M. lateralis* and the Caparaó population of this species (0.324 and 1.147 individuals per generation from *M. lateralis* to Caparaó and from Caparaó to *M. lateralis*, respectively), potentially reflecting the more isolated and restricted distribution of the Caparaó population (Table 2).

## Discussion

### The dual role of AF mountains: population divergence and persistence

Mountains may function both as generators and maintainers of diversity (Fjeldsa et al. 2012), and here we show a Atlantic Forest organisms that support this idea. First, the ancient AF highlands may generate diversity by fostering population isolation and divergence, as indicated by our population structure estimates with three distinct groups associated with three isolated mountain ranges. While geotectonic changes could explain phylogeographic breaks in the AF (e.g. Thomé et al. 2010, Amaro et al. 2012, Amaral et al. 2013), the shallow divergences found here cannot be reconciled with the major uplift events of eastern Brazil highlands, which very likely took place long before the estimated recent divergence times (< 20 k years). In addition, support for a model of synchronous isolation among the three populations is especially compelling in terms of large scale effects of climate across the entire range of subtropical AF. Interestingly, the strong population structure found here contradicts previous studies on other co-distributed montane organisms, which did not recover recent (< 100 k years) phylogeographic breaks (Amaro et al. 2012, Batalha-Filho et al. 2012, Peres et al. 2015). These findings suggest that although warbling-finches are forest-associated species that often use forest edges, such ecological flexibility was presumably not sufficient to preclude isolation and divergence. In contrast, the White-browed Warbler (*Myiothlypis leucoblephara*), for example, which is sympatric - and often syntopic - to *M*. *cabanisi* and *M*. *lateralis*, did not show detectable population structure (Batalha-Filho et al. 2012). These phylogeographic differences may be due to the interaction of traits and historical processes (see Zamudio et al. 2016 for a review), differences in altitudinal distribution, lack of power of Sanger datasets to detect shallow divergences (Amaral et al. 2018) or a combination of those factors, and warrants additional studies on both novel and previously studied organisms using subgenomic or genomic datasets.

Second, it has been long suggested that the MAF (Brown & Ab’Saber, 1979; Brown, 1987) and other Neotropical mountains (Fjeldså et al. 1999; García-Moreno & Fjeldså, 2000, Mastretta-Yanes et al. 2018) are safe harbors (i.e refuges) for montane and non-montane organisms during historical periods of harsh climates. Mountains may buffer population size changes of montane forest organisms during historical periods of unsuitable climate by holding moisture in the leeward slopes (Brown & Ab’Saber 1979) and by allowing shifts in elevational distributions, a process that may be even seen in ecological time (Moritz et al. 2008). In line with this observation, our estimates of historical demography echo the lack of strong population size fluctuations found in other co-distributed organisms (Amaro et al. 2012, Batalha-Filho et al. 2012, Peres et al. 2015, Pie et al. 2018), what underscores the notion that montane habitats are important refuges, hampering extinction of isolated population during periods of climate change.

While our results illustrate how montane habitats may act both as drivers and keepers of diversity, partial concordance with previous studies, specially in terms of population structure, is intriguing. New comparative phylogenomic studies will help understand the generality of the patterns found here, and contribute to tease apart the effects of history, ecology and issues related to number of loci. We hypothesize that population structure among highland species in the AF will be more common than previously thought, and in many cases may be detectable only using sufficiently sized nuclear datasets (e.g. Pie et al. 2018; present study). Fine-scale inferences of population structure will be essential not only to better describe patterns and infer processes involved in Neotropical montane diversification, but may also reveal important hotspots of divergence and cryptic diversity of conservation importance, which may have been overlooked due to recent divergences (as in the Araripe Highlands, Amaral et al. 2018).

### Historical climate changes explain diversification in MAF warbling-finches

A main question in the MAF system is whether one can disentangle the relative influence of topography and climate when explaining diversification patterns. For example, under a strict geomorphological model of diversification, genetic divergences among warbling finch populations should be coincident with the known major pulses of uplift of eastern Brazilian highlands during the last ca.100 mya (e.g. Melo et al. 1985, Meisling et al. 2001, Franco-Magalhães et al. 2014). However, we argue that one cannot rule out biogeographic scenarios in which historical climate changes taking place during or subsequent to the major uplift events had a main role on biotic diversification. In fact, the interplay between topography and climate implies that the expected model of diversification for warbling finches and other MAF organisms would be one of isolation and divergence on distinct highlands during the warm-interglacial periods, when populations retracted upwards over the mountains. In addition, any event of genetic admixture (i.e. introgression) between resulting lineages ensued when populations expanded downwards, during cold-glacial periods. The results of our coalescent analyses and the current altitudinal distribution of warbling finches confirm key components of this model. Specifically, the recent estimates of genetic divergence and past admixture (i.e. for the late Pleistocene) as well as their ocurrence on highlands during the present-day warm-interglacial period undoubtedly implicate a strong influence of historical climate changes on range shifts and associated population evolutionary dynamics across a longstanding rugged landscape. One especially intriguing result is the lack of recent variation in effective size despite signs of introgression. We hypothetize that secondary contact may not necessarily involve large population size fluctuation, as shifts in elevational distribution could occur without significant changes in effective size. In addition, small elevational shifts (e.g. only 100 m) could readly connect the two species (Fig. 1). Alternatively, we can also speculate that past gene flow may have erased signs of population expansion and contraction, or limited expansion associated with long-distance dispersal could explain introgression without large Ne variation.

### Elevational distribution and its role in population isolation

Population structure was detected in the northern species (*M. lateralis*) but was absent in the southern species (*M. cabanisi*), and it is possible that the phylogeographic structure of *Microspingus* warblers may reflect historical differences in forest dynamics between southeastern and southern AF. One interesting biological difference between *M. cabanisi* and *M. lateralis* is their distinct elevational range: populations of *M. lateralis* are found only above 900 m, while in the southernmost part of its range, *M. cabanisi* reaches sea-level. This pattern may be a consequence of either thermal niche divergence (Janzen 1967), biological interactions (i.e. competition) or simply a latitudinal compensation of elevational distribution (Barry 1992). In any case, differences in altitudinal ranges across a latitudinal gradient may lead to greater opportunities for isolation in the populations in the northern extreme edges of SE/S subtropical habitats than those in the southern edges. Additional multi-taxon studies are warranted to test this hypothesis, as it could explain distinct levels of endemism in different portions of the MAF.

### Caparaó highlands as a hotspot of divergence

The Caparaó is the highest mountain massif in the AF, reaching ca. 2900 m asl. Despite its small area compared to the neighbouring highlands (Fig. 1), this mountain system not only harbors a strongly differentiated population of *Microspingus* warbling-finches, but also endemic taxa (*i.e. Caparaonia* lizards, Rodrigues et al. 2009) whose divergences from taxa in other larger AF highlands are possibly much older than our divergence estimates. This suggests that the Caparó highlands possibly represent a microrefugium (*sensu* Rull 2009) with a long-term history of connection and isolation with the neighboring highlands. This makes the Caparaó highlands an overlooked hotspot of population divergence and cryptic diversity, similarly to the Chapada Diamantina highlands (Amaral et al. 2013) and the Araripe plateau (Amaral et al. 2018). The presence of both recent and old endemic lineages in the Caparaó mountains may be related to higher population persistence in contrast to lower neighboring mountains. Interestingly, although the Caparáo population of *M. lateralis* diverged from the remaining *M. lateralis* populations, genetic differentiation was not detected between the other two currently isolated (by the warm lowlands of the Paraíba do Sul river valley) Serra do Mar and Serra da Mantiqueira populations. This could be related to a greater geographic proximity between the latter, thus allowing genetic homogeinization. Given that population size changes may be buffered by shifts in elevational ranges and that extreme interglacial-warm periods could crash mountain-top populations (Moritz et al. 2008), it is possible that higher mountain ranges are related to higher persistence, higher levels of endemicity, wider temporal ranges of population divergence compared to lower neighboring mountain ranges. Populations of many species that occur in Mantiqueira and Serra do Mar mountain systems can also be found in Caparaó and offer exciting future opportunities to identify cryptic diversity.

### The São Paulo and Caparaó subtropical gaps

Low passes and inter-mountain valleys often constitute formidable barriers to dispersal in montane tropical organisms (e.g. Africa - Bowie et al. 2006; Andes - Cadena et al. 2007, Winger and Bates 2015). However, such ecological barriers to genetic exchange remain largely overlooked in the MAF biota. In our study, more than 100 kilometers of lowlands that reach elevations down to 250 m a.s.l. in the Ribeira de Iguape River Valley (Karl et al. 2013) separate *M. lateralis* from *M. cabanisi*. This coincides with range and phenotypic discontinuities in other cold-adapted avian species complexes, including passerines (e.g. *Hemitriccus* flycatchers, *C. thoracica* warbling-finch) and non-passerines (*Stephanoxis* hummingbirds, Cavarzere et al. 2014), as well as with a narrow zone of secondary contact between cold-adapted lineages of *Bombus* bees (Françoso et al. 2016). Lower elevation forests also seem to isolate the highland biota of Caparaó from the ones of southern mountain ranges. We suggest that these lower elevation forests represent the ecological barrier underlying the differentiation in these and possibly other MAF organisms and could be named “São Paulo subtropical gap” and “Caparaó subtropical gap”. These ecological gaps highlights the influence of historical climate changes and associated shifts in the geographic distribution of MAF organisms, and coincides with the genetic differentiation observed in our study system. Additional studies will show if the historical processes affecting montane organisms in this region are shared, thus generating population divergence in other co-distributed organisms.

## Acknowledgments

We thank C. Assis, A. Bianco, B. Genevcius, G. Macedo, D. N. Oliveira, V. Q. Piacentini, R. Quagliano, C. F. Schwertner, J. Vitto for their excellent field assistance. Michael Sovic shared fastsimcoal models. Special thanks to B. Faircloth for providing extensive help with UCE processing and SNP calling. ICMBio provided collection permits (14673, 30829, 30835, 30836, 30837, 30838, 30840, 30841, 30844, 37011, 39935, and 47205) and access to National Parks (P. N. Caparaó., P. N. Serra da Bocaina, P. N. São Joaquim, P. N. Serra Geral, P. N. Aparados da Serra). This research has been evaluated by UNIFESP Ethics Commitee (CEP 0069/12). D. N. Oliveira helped with lab work. We also thank private landowners for access to land under their care. Financial support was provided by FAPESP and the BIOTA program (2011/50143-7, 2011/23155-4, 2015/18287-0, 2014/00113-2, 2015/12551-7, 2017/25720-7, 2018/17869-3), a joint NSF, FAPESP and NASA grant (Dimensions US-Biota-SP, FAPESP 2013/50297-0 and DOB 1343578,), CNPq and Coordenação de Aperfeiçoamento de Pessoal de Nível Superior - Brasil (CAPES) - Finance Code 001. For helpful discussions we thank Scott V. Edwards, J. M. B. Alexandrino, C. Assis, Fernando D’Horta, V. Q. Piacentini, Cibele Biondo, Camila Marques, Guilherme Cavicchioli, Marcel Neves, Deborah Nacer, Bruna Gimenez, Estela Silva and the FAPESP/NSF AF-Biota group.

## Supplementary Information

**Table S1.**
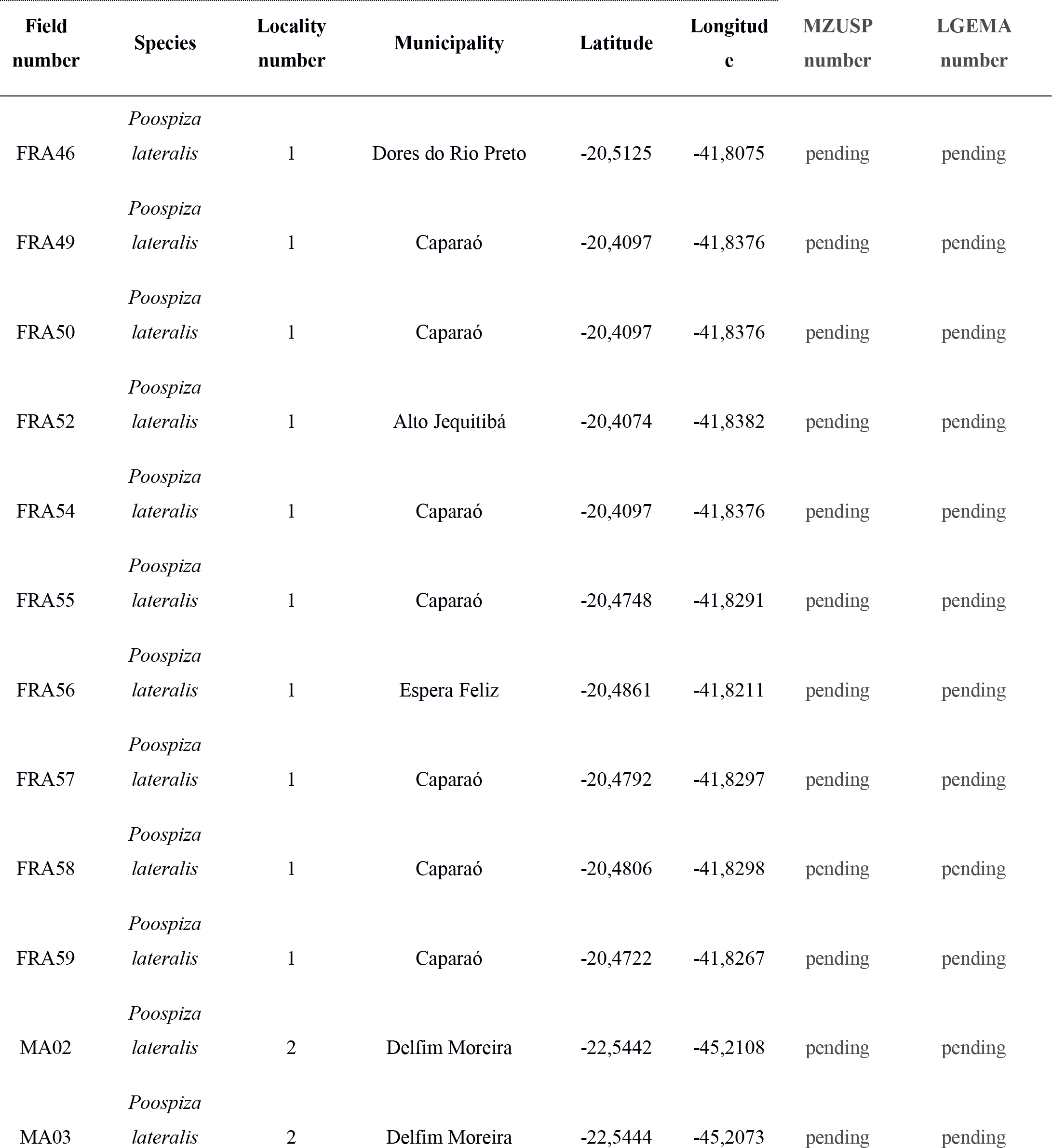
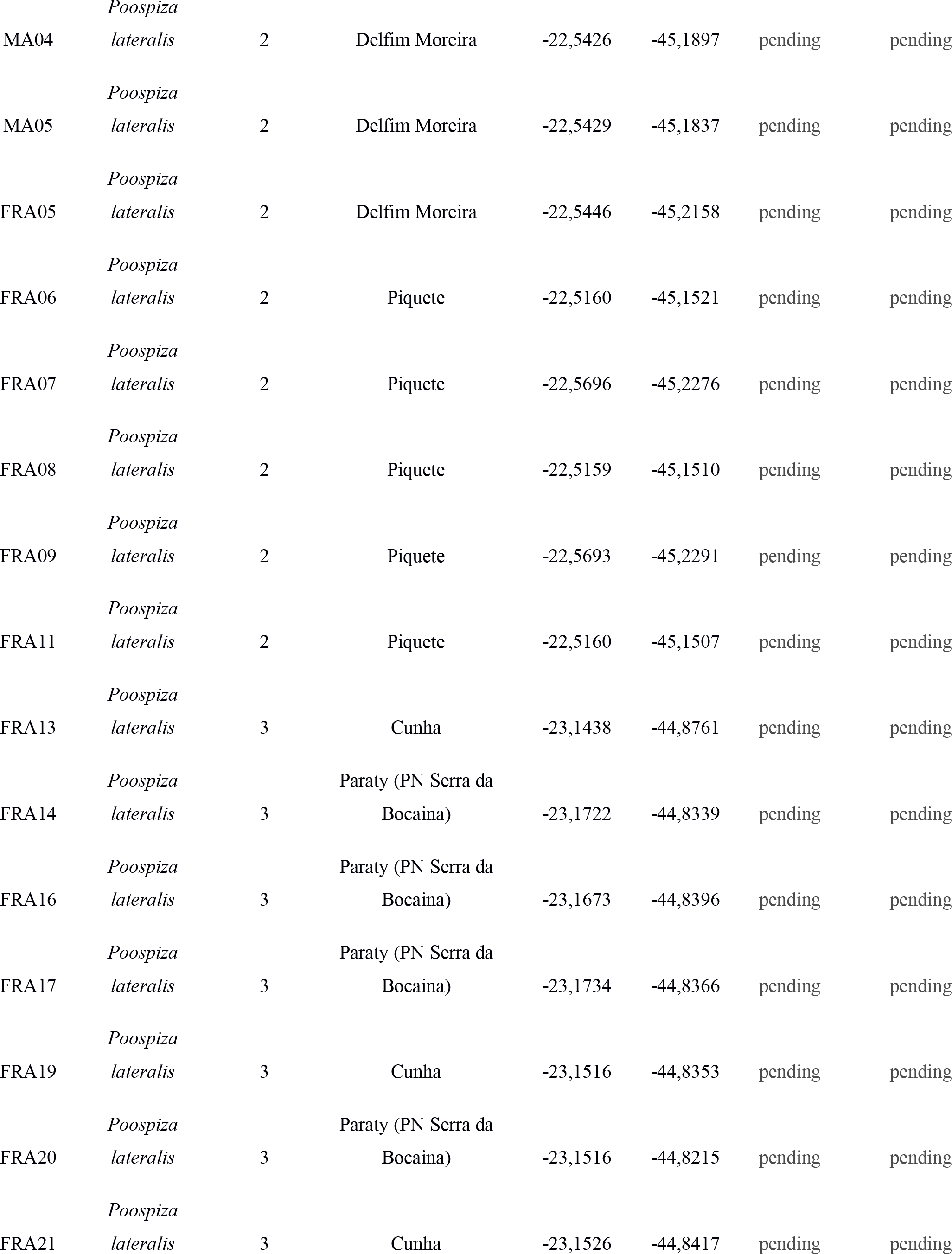
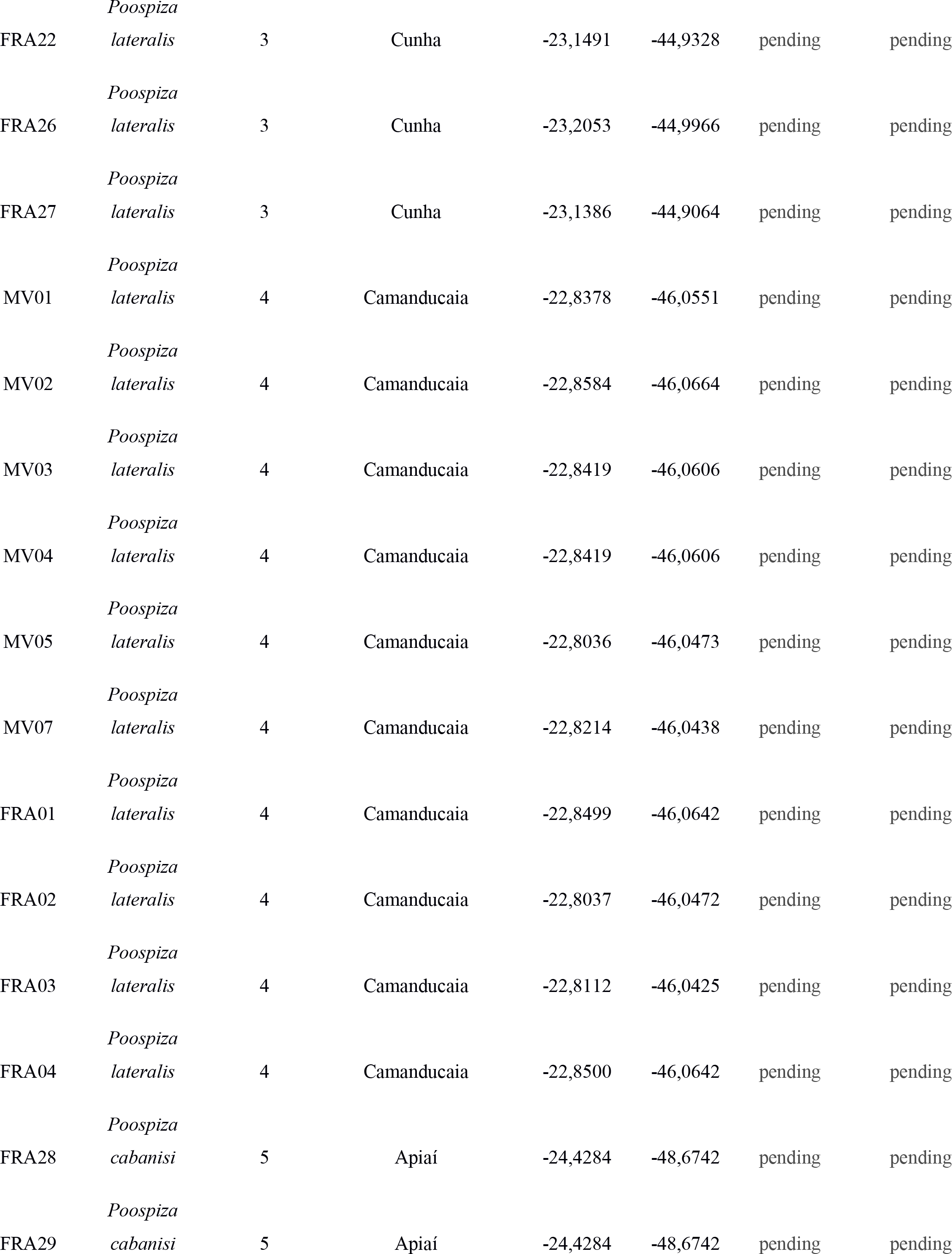
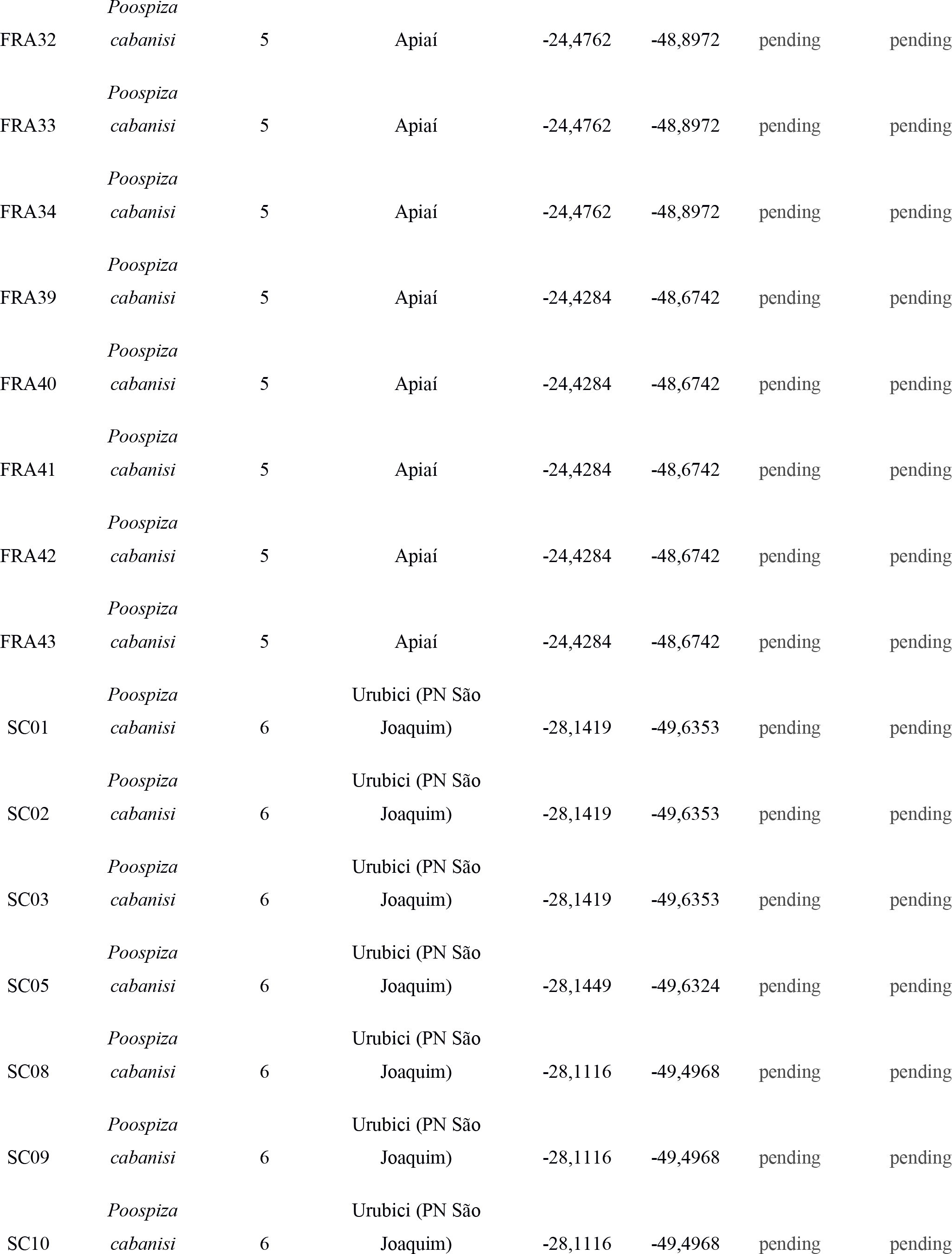
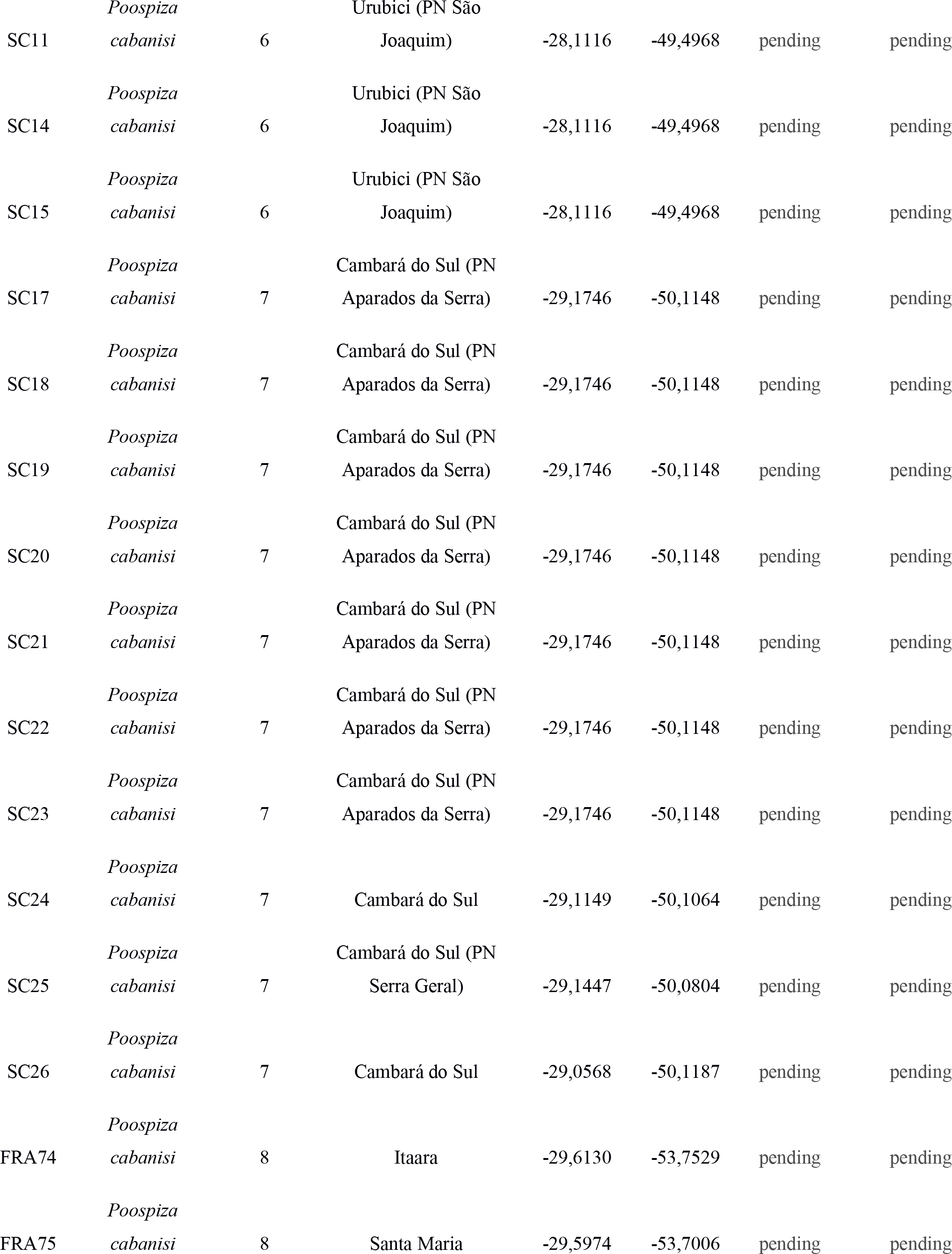
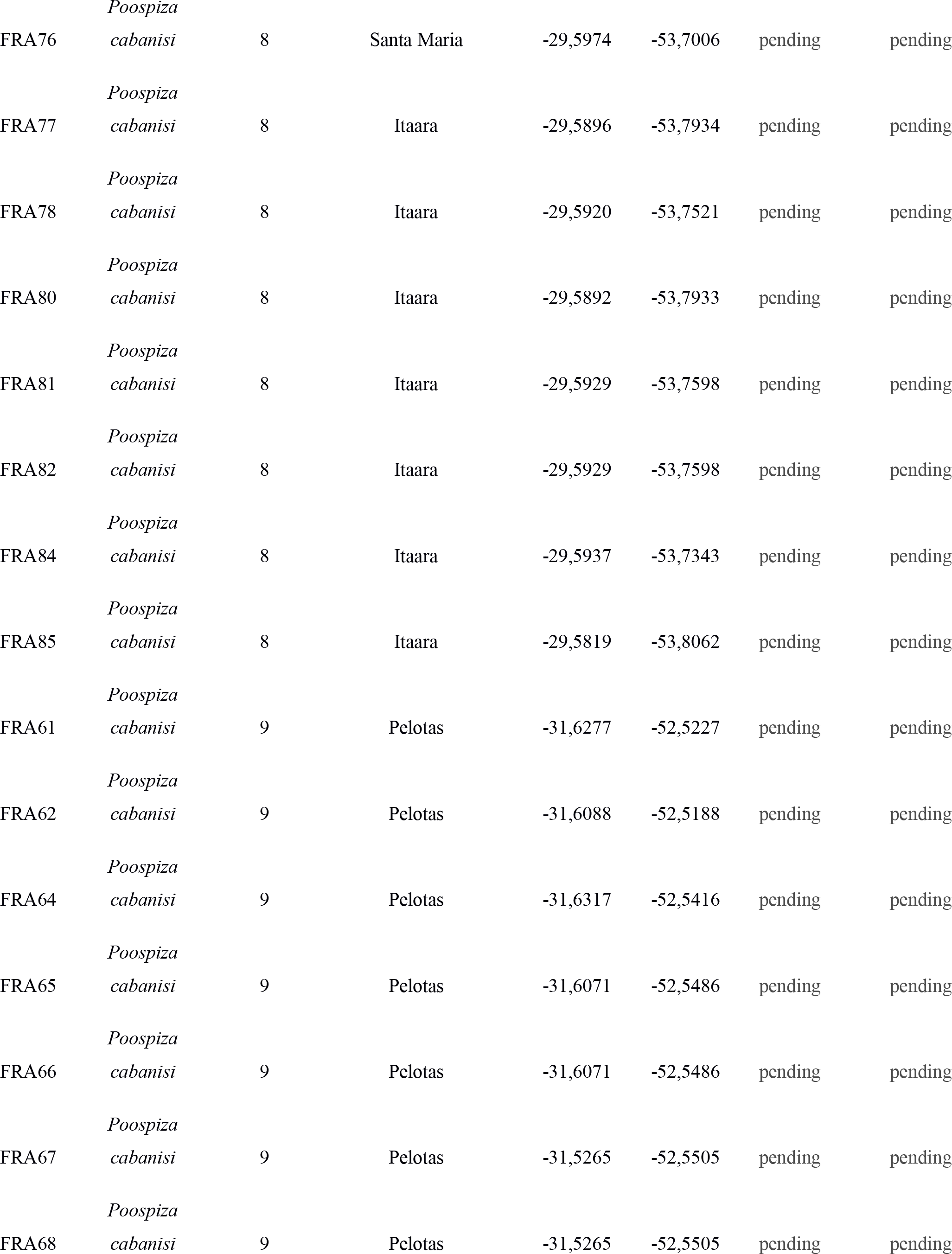
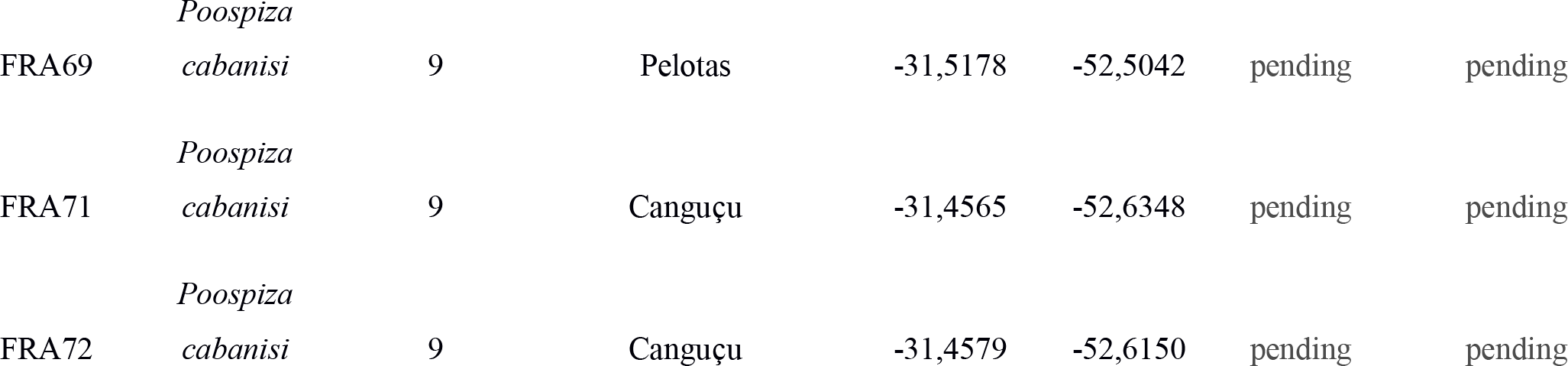
Samples used in the present study.

**Figure S1.**
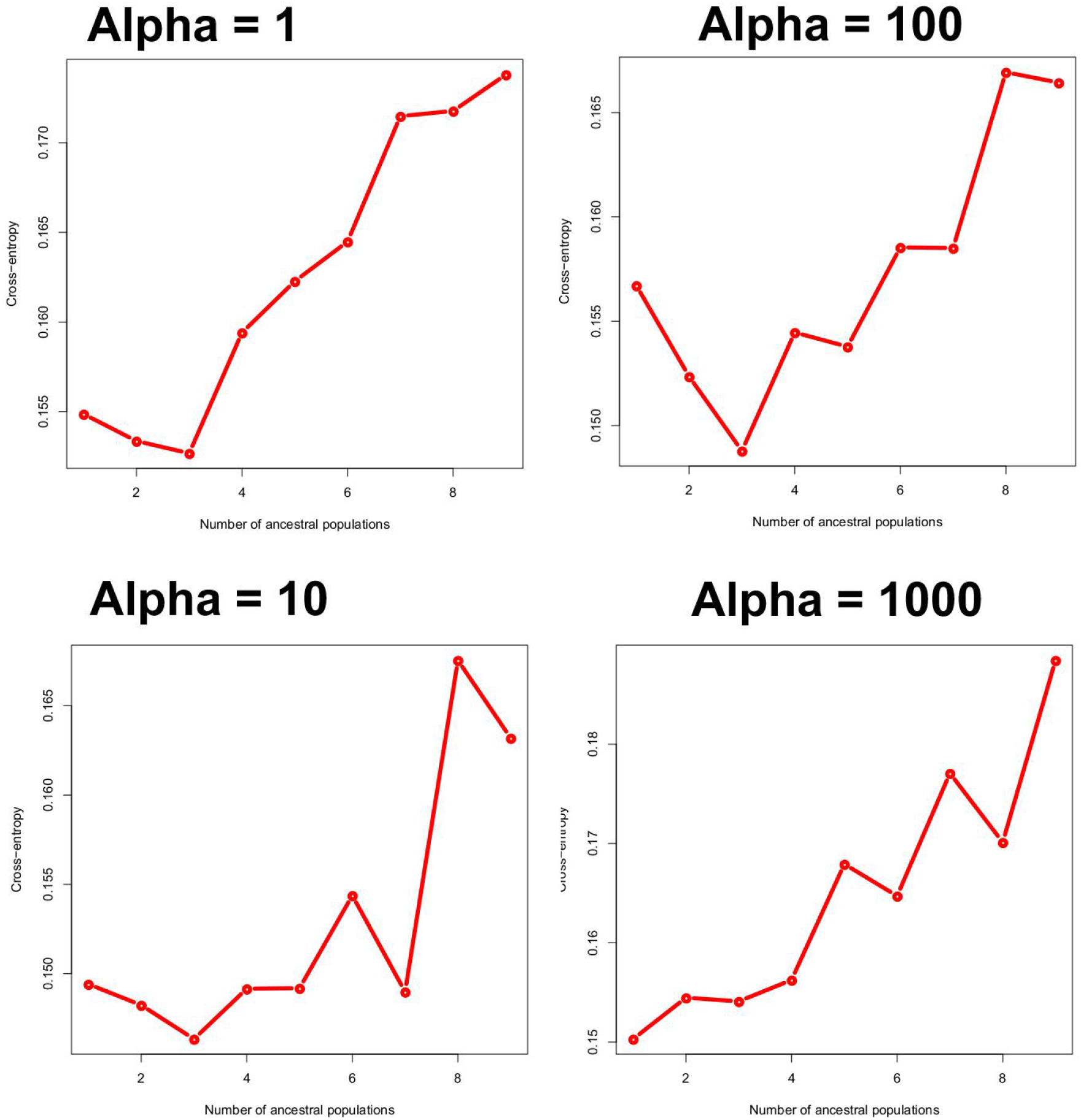
Cross-entropy sNMF runs for alpha values of 1, 10, 100 and 1,000.

**Figure S2.**
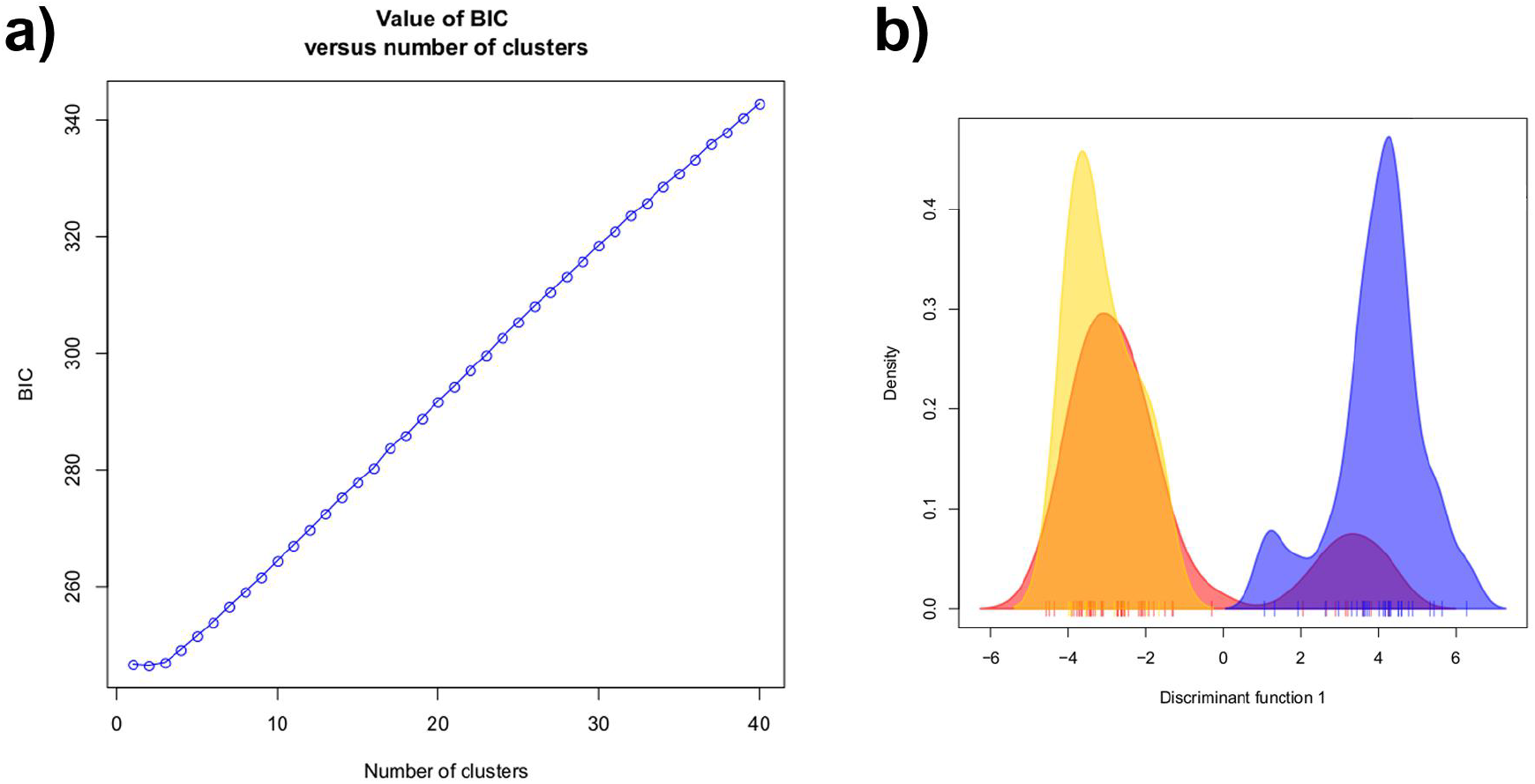
BIC values of (a) K-clustering method and (b) DAPC plot for two populations. Colors in the DAPC plot follow the ones assumed for Figures 1 and 2.

## Data Accessibility

Dryad XXXX and SRA XXXX

Raw Illumina files, Set of SNPs, Joint SFS

## Author contributions

FRA, DFA and GT designed the study. FRA, JACM and GT performed field work. FRA, DFAS and GT performed data analysis. FRA, KCMP, CYM and MJH contributed funds for lab or field work. All authors participated in the discussion of the results. FRA, MMC and GT wrote the manuscript with input from the other authors. All authors approved the final version of the manuscript.

